# Forward time-series causal inference based on forest weights and conditional dependence

**DOI:** 10.64898/2026.10.02.756132

**Authors:** Shigeki Kishi

**Affiliations:** Research Center for Agricultural Information Technology, National Agriculture and Food Research Organization, 1-13-1 Kannondai, Tsukuba, Ibaraki 305-0856, Japan

## Abstract

Unveiling directional causality in complex, high-dimensional dynamical systems remains a fundamental challenge. Conventional nonparametric approaches, such as Convergent Cross Mapping, require computationally intensive surrogate tests and frequently yield false-positive detections under strong coupling due to generalized synchronization. Here, we present F-CODEC (Forest-weighted Conditional Dependence Coefficient), a forward-looking causal discovery framework integrating generalized random forests with adaptive rank conditioning. Nonparametrically conditioning on the target variable’s intrinsic delay coordinates systematically neutralizes shared manifold representations, suppressing false positives without surrogate data. Furthermore, an analytical time-series threshold allows instant, rigorous hypothesis testing directly from raw observations. We demonstrate F-CODEC’s resilience against generalized synchronization and high-dimensional noise across canonical chaotic benchmarks and multi-species network motifs. Applying F-CODEC to historical lynx–hare cycles resolves delayed trophic forcing, while in-situ IoT honeybee monitoring captures real-time microclimate-acoustic causal surges preceding colony-level behavioral events. F-CODEC establishes a practical, scalable, and surrogate-free tool for causal discovery in nonlinear dynamical systems.

## Introduction

Real-world dynamical systems inherently exhibit complex behaviors, characterized by pervasive nonlinearities, delayed feedback loops, multiscale temporal dependencies, threshold responses, and state-dependent interactions ^1,2^. The rapid proliferation of automated sensor arrays and continuous monitoring technologies has enabled the high-resolution capture of these dynamics, yielding an unprecedented accumulation of time-series data across diverse disciplines, including ecology, climatology, neuroscience, and industrial engineering. In such intricate, high-dimensional settings, extracting genuine causal architectures from observational data is paramount for resolving underlying governing mechanisms, anticipating critical state transitions, and designing effective, targeted interventions.

To detect causality within nonlinear dynamical systems, Convergent Cross Mapping (CCM) ^3^ has emerged as a cornerstone methodology. Grounded in Takens’ delay embedding theorem ^4^, CCM operates on a “backward-looking” state-space reconstruction principle: it posits that if a causal variable *X* forces an effect variable *Y*, the historical trajectory of *X* is necessarily encoded in the state space of *Y*, allowing the reconstructed shadow manifold MY to cross-predict the historical states of *X* (*M_X_*). This foundational paradigm has catalyzed extensive empirical breakthroughs and spurred the continuous development of the Empirical Dynamic Modeling (EDM) framework ^5^. A similar backward-looking topological logic also underpins other nonlinear causal measures, such as recurrence-based quantification and the joint distribution of inter-point distances ^6^.

Despite its widespread adoption, state-space causal inference faces fundamental theoretical and computational bottlenecks ^7,8^. Chief among these is the vulnerability to generalized synchronization under strong dynamical coupling ^9–11^. When coupling strength exceeds a moderate threshold, the attractor manifolds of interacting variables collapse into a mutually diffeomorphic geometry. Under this synchronized regime, backward cross-mapping succeeds symmetrically in both directions, erroneously identifying strong bidirectional causation in strictly unidirectional systems ^12^. Consequently, state-space methods are notoriously prone to severe false-positive detections in strongly interacting regimes. Furthermore, establishing statistical significance in these frameworks typically necessitates benchmarking against thousands of surrogate time series to preserve minimal temporal characteristics such as autocorrelation ^13,14^. This surrogate-testing procedure imposes severe computational costs that scale poorly with time-series length and network dimensionality, fundamentally impeding real-time and high-throughput causal discovery. While forward-looking information-theoretic frameworks such as Transfer Entropy circumvent backward cross-mapping, they likewise remain tethered to computationally intensive surrogate testing ^15,16^.

To overcome these challenges, model-free conditional dependence measures offer a promising alternative. Recently, Azadkia and Chatterjee introduced the conditional dependence coefficient (CODEC), which rigorously quantifies conditional dependence without distributional or parametric assumptions ^17^. Grounded in rank-based topological proximity, CODEC assesses whether observations that are geometric nearest neighbors in the joint space (*X*, *Z*) exhibit closer ranks in the target variable *Y* than those that are nearest neighbors solely in the conditioning space *Z*, thereby measuring the unique predictive information contributed by *X* beyond *Z*. Although CODEC was originally developed for static feature selection under the assumption of independent and identically distributed (i.i.d.) data, its forward conditional formulation provides an ideal foundation for dynamical systems. However, standard CODEC suffers from severe limitations when applied to time-series and dynamical systems. Because it relies on simple Euclidean 1-nearest-neighbor searches across the conditioning manifold, standard CODEC is highly vulnerable to observational noise and uninformative dimensions. As the conditioning dimension increases, particularly when the dimensions are noisy and less informative, standard nearest-neighbor matching rapidly succumbs to the curse of dimensionality, leading to severe information loss, distorted neighbor distances, and deteriorated statistical power.

This limitation can be resolved by replacing rigid Euclidean distances with adaptive tree-based similarity metrics. Generalized Random Forests (GRF) ^18^ construct dynamic kernel weights, termed forest weights, based on the frequency with which observation pairs co-occur in the terminal leaves of an ensemble of decision trees. By recursively partitioning state space along the most informative coordinates, forest weights automatically adapt to local manifold geometry, effectively filtering out irrelevant noise dimensions and substantially mitigating the curse of dimensionality inherent to high-dimensional Euclidean nearest-neighbor searches ^19^.

Here, we propose the Forest-weighted Conditional Dependence Coefficient (F-CODEC), a forward-looking, model-free causal inference framework that integrates generalized random forests with a modified conditional dependence metric for nonlinear dynamical systems. Instead of backward cross-mapping, F-CODEC operates strictly forward in time, quantifying the pure conditional predictive contribution of a lagged causal candidate *X_t-L_* on the future target state *Y_t_*_+1_, nonparametrically conditioned on the target’s own intrinsic delay-coordinate history **Z**_cond_ = [*Y_t_*, *Y_t_*_–1_, …, *Y_t–E_*_+1_]. Crucially, by nonparametrically conditioning on the target’s autonomous delay coordinates and candidate confounders, F-CODEC projects out intrinsic dynamical persistence and systematically removes dependencies attributable to observed conditioning variables, thereby suppressing spurious causal signals from generalized synchronization and transitive pathways under the assumption of causal sufficiency. Furthermore, by formalizing the manifold contraction rate via rank-space multiple determination (*R_rank_*^2^), we derive an analytical, time-series-corrected significance threshold (*T*_crit_TS_) that establishes rigorous, instant hypothesis testing directly from raw observations.

In this study, we systematically evaluate the capability of F-CODEC across synthetic dynamical models and real-world empirical benchmarks. First, we test the framework on coupled chaotic logistic map systems across varying coupling strengths to demonstrate its resilience against generalized synchronization, compared to CCM. Second, we examine three-species network motifs (cascade chains and common-cause fan-outs) to evaluate the systematic pruning of indirect and spurious linkages. Third, we apply F-CODEC to historical Canada lynx and snowshoe hare population cycles to resolve directional trophic causality and characteristic multi-year time lags. Finally, we implement the framework on continuous in-situ IoT streaming data from honeybee (*Apis mellifera*) hives, tracking time-varying causal strengths and Jacobian interactions between microclimate fluctuations and collective acoustic responses during colony-level behavioral events.

## Results

### Quantitative Metric Scaling and Synchronization Resilience in Coupled Chaos

To evaluate how the F-CODEC statistic responds to coupling strength and whether it is susceptible to generalized synchronization, we evaluated unidirectional coupled logistic maps (*X* → *Y*) across coupling strengths *C* ∈ [0.0,0.5] (Fig. 1). In the true causal direction (*X* → *Y*), the mean conditional dependence coefficient *T_n_* increased monotonically from *T_n_* = 0.004 (95% CI: [-0.060,0.068]) at *C* = 0.0 to *T_n_* = 0.163 at C = 0.1, reaching *T_n_* = 0.405 at *C* = 0.5 (Fig. 1). For all coupled regimes (*C* ≥ 0.1), *T_n_* decisively exceeded the asymptotic analytical 95% threshold (*T*_crit_ = 0.060). Conversely, in the uncoupled reverse direction (*Y* → *X*), *T_n_* remained constrained near zero across all coupling strengths (*T_n_* ≤ 0.006, 95% CI upper bound ≤ 0.017), remaining well below *T*_crit_ (Fig. 1). This confirms that nonparametrically conditioning on the target’s intrinsic delay coordinates systematically neutralizes the shared manifold geometry induced by generalized synchronization.

**Figure 1.**
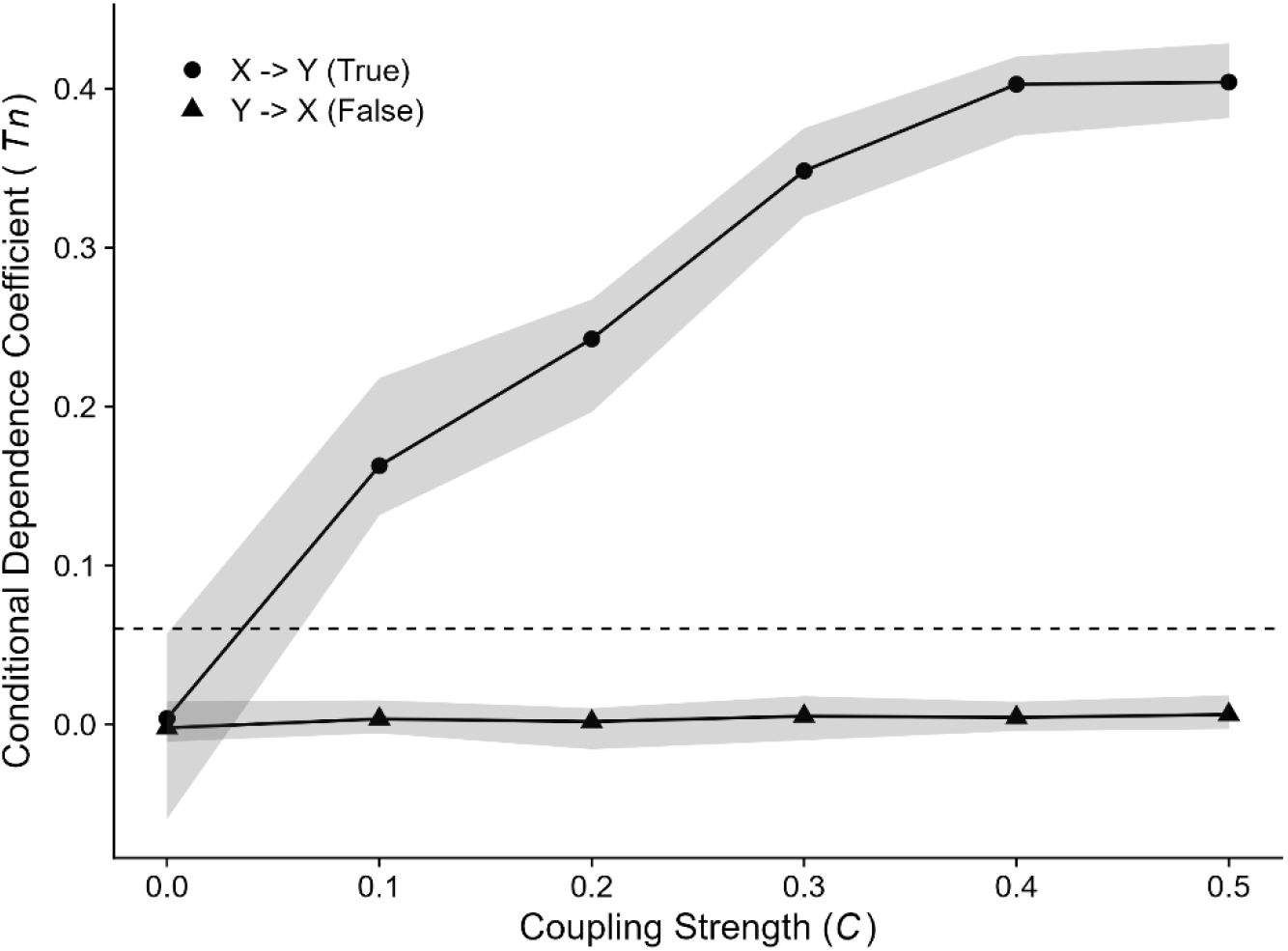
Mean conditional dependence coefficient (*Tn*) evaluated for the true causal link (*X* → *Y*, filled circles) and the spurious reverse link (*Y* → *X*, filled triangles) in a unidirectionally coupled chaotic logistic map system across coupling strengths *C* from 0.0 to 0.5. Shaded gray ribbons represent empirical 95% percentile confidence intervals (2.5th to 97.5th percentiles) across 15 independent simulation trials. The horizontal dashed line denotes the analytical asymptotic 95% significance threshold under the null hypothesis (*T*_crit_ = 0.060 for *N*_eff_ = 299). In the true causal direction, *Tn* increases monotonically with coupling strength and exceeds the critical threshold for *C* ≥ 0.1. In the uncoupled reverse direction, *Tn* remains strictly constrained near zero and below the threshold across all coupling regimes, demonstrating complete resilience against generalized synchronization artifacts.

We next benchmarked causal detection sensitivity and false-positive control under high-dimensional noise against alternative methodologies across C ∈ [0.0,0.7] (Fig. 2). In the absence of uninformative noise dimensions, both F-CODEC and classical Euclidean Raw CODEC achieved 100% true-positive detection for *C* ≥ 0.1 while maintaining nominal 0% false-positive rates (Fig. 2a, b). However, upon adding five uninformative noise lag dimensions (**Z**_cond_ ∈ R^7^), Raw CODEC suffered severe statistical power degradation due to distance concentration, with detection rates dropping to 16% at *C* = 0.1 and 50% at *C* = 0.2 (Fig. 2d). In contrast, F-CODEC maintained 100% true-positive sensitivity and 0% false positives across all non-zero coupling strengths (Fig. 2c). Comparison with alternative frameworks highlighted key limitations of existing methods: FORCC exhibited slight false-positive inflation in the reverse direction under strong coupling (*C* ≥ 0.6, detection rate reaching 26.5%) (Fig. 2e), whereas conventional CCM suffered severe false-positive inflation in the uncoupled reverse link (*Y* → *X*), escalating monotonically from 12% at *C* = 0.1 to 90% at *C* = 0.7 as generalized synchronization emerged (Fig. 2f).

**Figure 2.**
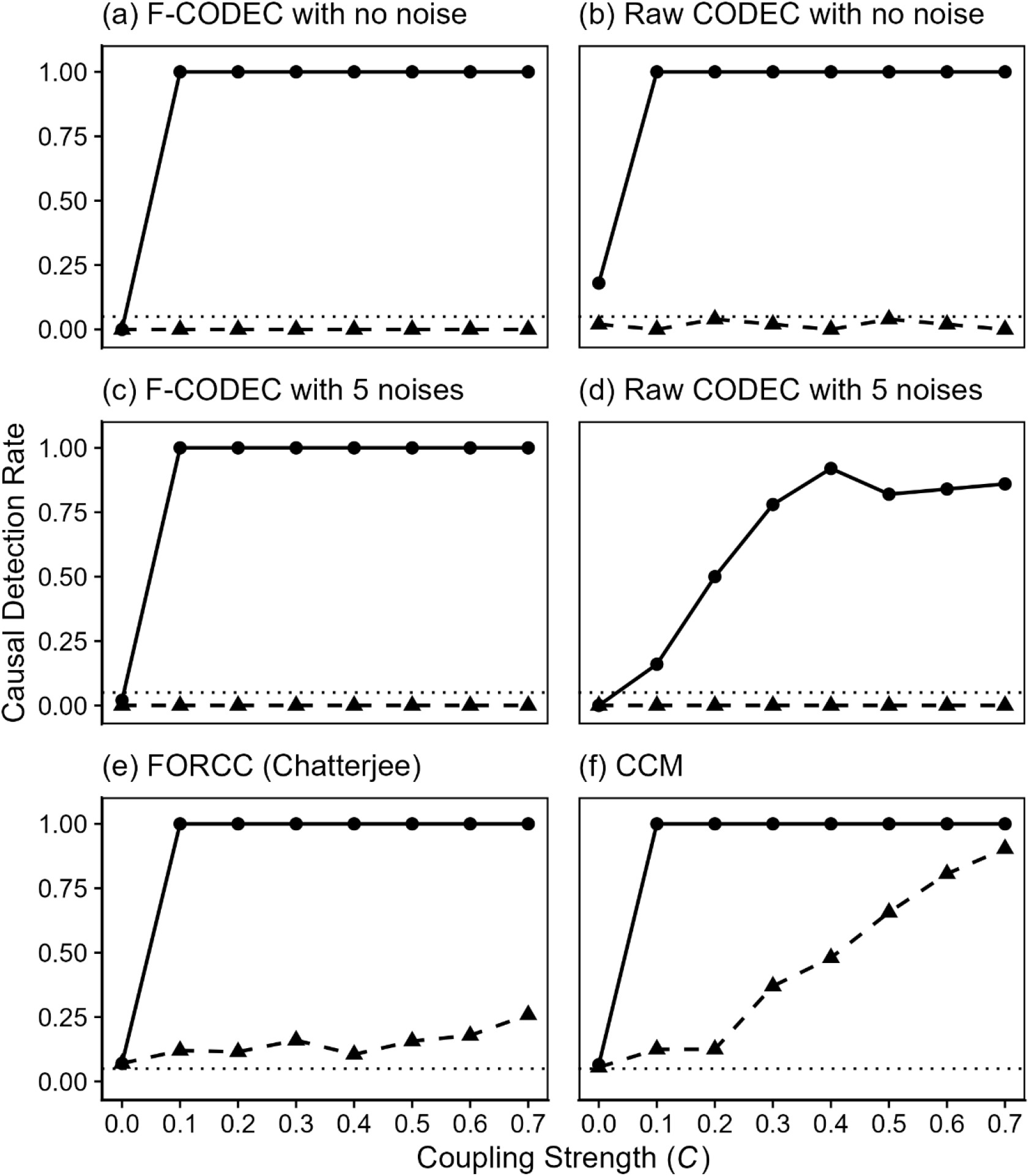
Systematic benchmark of causal detection performance under varying coupling strengths and high-dimensional noise. Causal detection rates at nominal *α* = 0.05 for the true causal direction (*X* causes *Y*, solid lines with filled circles) and the uncoupled reverse direction (*Y* causes *X*, dashed lines with filled triangles) evaluated across coupling strengths *C* from 0.0 to 0.7. Horizontal dotted lines denote the nominal 5% false-positive baseline. (a) F-CODEC without uninformative noise dimensions (**Z**_cond_ in R^2^). (b) Raw unweighted Euclidean CODEC without noise dimensions (**Z**_cond_ in R^2^). (c) F-CODEC under 5-dimensional uninformative noise confounding (**Z**_cond_ in R^7^), maintaining 100% true-positive sensitivity while keeping false positives at zero. (d) Raw unweighted Euclidean CODEC under 5-dimensional noise confounding (**Z**_cond_ in R^7^), showing severe statistical power degradation caused by the curse of dimensionality. (e) FORCC (Forest-weighted Chatterjee correlation), showing slight false-positive inflation under strong coupling (*C* ≥ 0.6). (f) Conventional Convergent Cross Mapping (CCM), exhibiting severe false-positive inflation (up to ∼90%) in the uncoupled reverse direction (*Y* → *X*) as generalized synchronization emerges at high coupling strengths. Results are based on 50 realization trials for a–d and 200 trials for e and f.

### Disentangling Direct Causality in Multi-Species Network Motifs

To verify whether F-CODEC distinguishes direct causation from transitive indirect cascades and common-cause confounding, we evaluated canonical three-species interaction motifs (Fig. 3). In the cascade chain (*X* → *Y* → *Z*), F-CODEC exclusively identified genuine direct interactions (*X* → *Y*: median *T_n_* = 0.290, IQR: [0.268, 0.301]; *Y* → *Z*: median *T_n_* = 0.338, IQR: [0.320, 0.358]), with all 50 realization trials exceeding the analytical threshold (*T*_crit_TS_ ≈ 0.047) (Fig. 3a). When testing the indirect transitive link (*X* → *Z*), conditioning on both the target history and intermediate mediator *Y* effectively eliminated the transitive artifact (median *T_n_* = –0.005, IQR: [–0.009, – 0.002]) (Fig. 3a). All non-causal reverse directions (*Y* → *X*, *Z* → *Y*, and *Z* → *X*) were suppressed below the threshold (median *T_n_* ≤ –0.003) (Fig. 3a).

**Figure 3.**
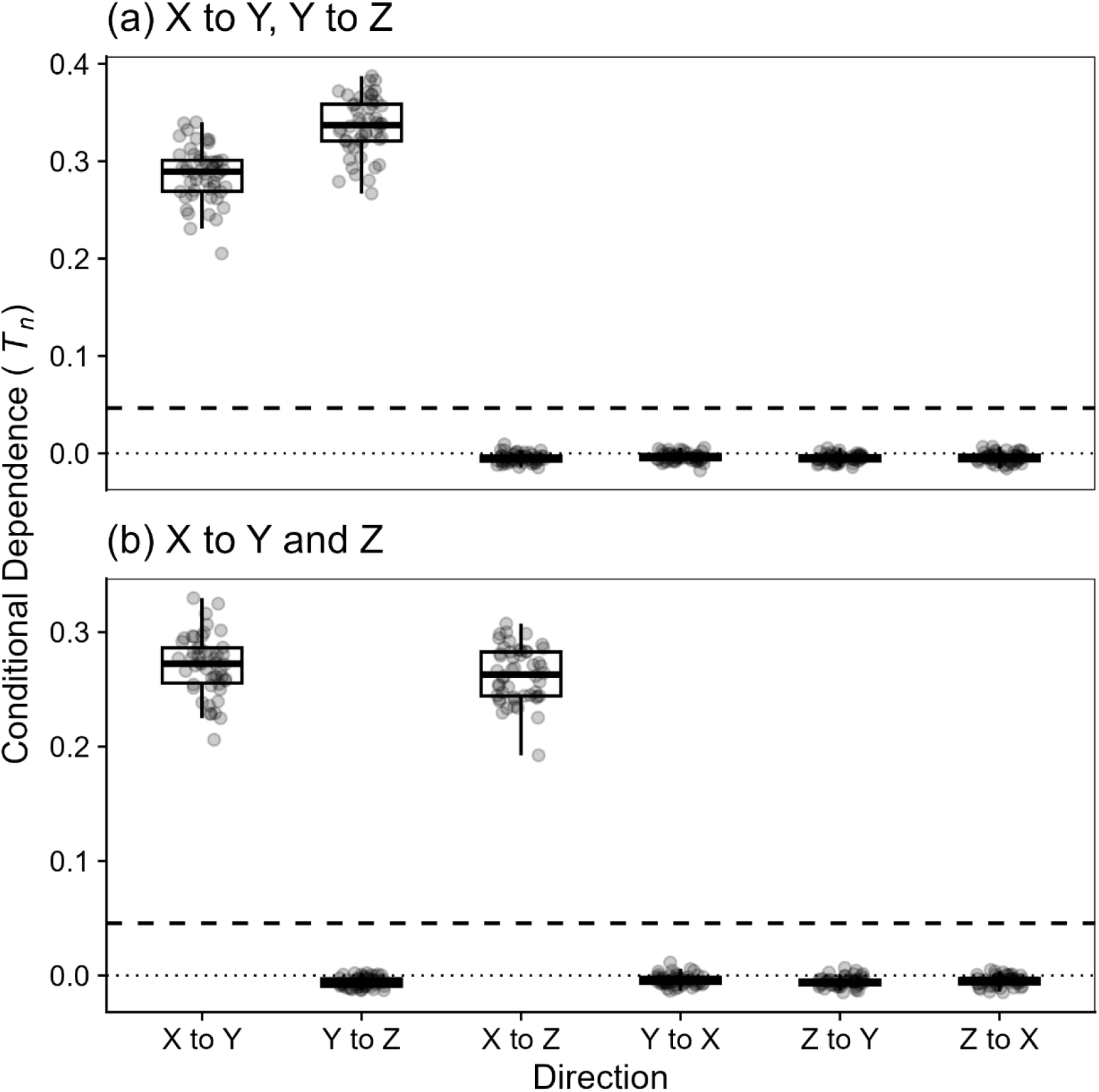
Systematic causal discovery and network pruning in synthetic three-species dynamical motifs. Boxplots and jittered points depict conditional dependence coefficients (*T_n_*) across 50 independent realization trials for all six directional permutations. Boxplot centerlines represent the median; lower and upper hinges indicate the first and third quartiles (IQR); whiskers extend to 1.5 × IQR. Horizontal dashed lines denote the trial-averaged analytical significance threshold (*T*_crit_TS_ ≈ 0.047, nominal *α* = 0.05); horizontal dotted lines indicate the zero baseline (*T_n_* = 0). (a) Multi-trophic cascade chain (*X* → *Y* → *Z*). F-CODEC exclusively identifies genuine direct causal links (*X* → *Y*, median *T_n_* ≈ 0.29; *Y* → *Z*, median *T_n_* ≈ 0.34), avoiding the indirect transitive artifact (X → Z) and suppresses all non-causal reverse directions below the analytical threshold. (b) Common-cause fan-out motif (*Y* ← *X* → *Z*). Direct causal emissions from driver *X* to downstream targets (*X* → *Y*, median *T_n_* ≈ 0.27; *X* → *Z*, median *T_n_* ≈ 0.26) are robustly detected, whereas conditioning on the true causal parent *X* eliminates spurious correlations between co-regulated variables (Y → Z and Z → Y, median *T_n_* ≈ 0).

Similarly, in the common-cause fan-out motif (*Y* ← *X* → *Z*), F-CODEC detected direct causal emissions from driver *X* to downstream targets (*X* → *Y*: median *T_n_* = 0.272, IQR: [0.256, 0.286]; *X* → *Z*: median *T_n_* = 0.263, IQR: [0.244, 0.282]) (Fig. 3b). By conditioning on the true causal parent *X*, the apparent spurious correlation between co-regulated variables (*Y* → *Z*) was resolved, yielding median *T_n_* values of –0.006 for *Y* → *Z* and –0.005 for *Z* → *Y* (Fig. 3b). Non-causal backward links (*Y* → *X* and *Z* → *X*) were suppressed near zero (median *T_n_* ≤ –0.004) (Fig. 3b).

### Resolving Asymmetric Trophic Causality in Canada Lynx and Snowshoe Hare Cycles

Applying F-CODEC to the 91-year historical pelt records of Canada lynx and snowshoe hare resolved clear asymmetric causal directionality and characteristic multi-year time lags (Fig. 4, Fig. 5). Autonomous embedding dimension scanning identified optimal dimensions of *E*\* = 2 for both species. Bidirectional forward lag scans (*L* ∈ [0, 8] years) revealed that the bottom-up direction (*Hare* → *Lynx*) yielded no statistically significant predictive gain beyond the lynx’s own historical persistence across all evaluated annual lags (*T_n_* ≤ –0.009, remaining below *T*_crit_TS_ ≈ 0.052) (Fig. 4a). In contrast, the top-down hypothesis (Lynx → Hare) identified significant predictive forcing peaking prominently at a 2-year delay (*T_n_* = 0.106, decisively exceeding the nominal threshold *T*_crit_TS_ = 0.080). Notably, this 2-year peak remains statistically robust even after accounting for multiple lag testing via conservative Bonferroni adjustment (*T_n_* > *T*_crit_Bonf_ ≈ 0.096), corroborating established ecological lag times required for predator numerical response and predation pressure to manifest in prey populations.

**Figure 4.**
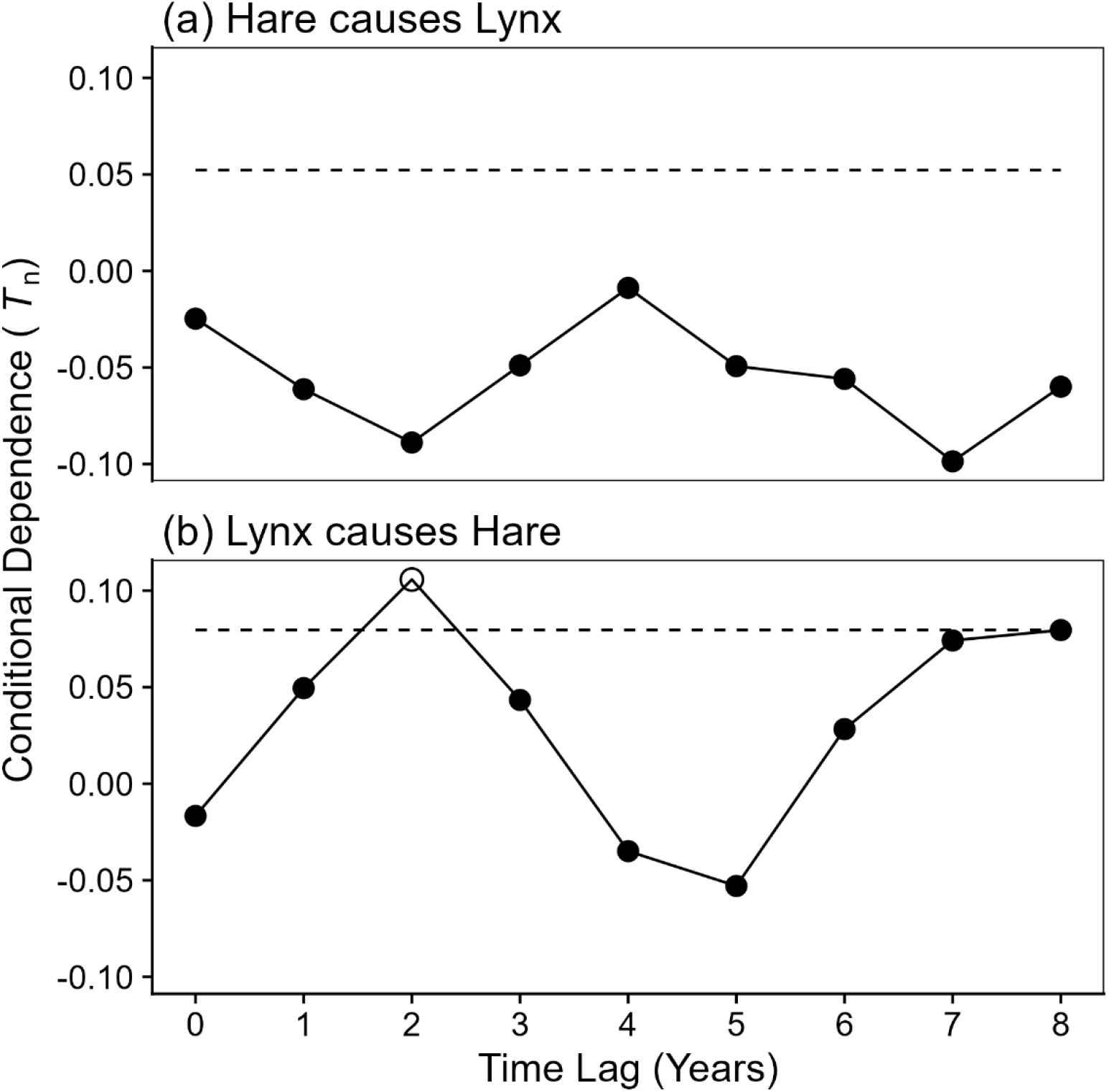
Bidirectional forward lag scan of causality in the Canada lynx and snowshoe hare ecological cycle. Conditional dependence coefficients (*T_n_*) evaluated across forward time lags ranging from 0 to 8 years. Open circles denote statistically significant causal linkages where *T_n_* exceeds the rank-space autocorrelation-corrected threshold (*T*_crit_TS_, horizontal dashed lines; nominal *α* = 0.05); filled circles represent non-significant lags. (a) Bottom-up direction (Hare → Lynx) and (b) Top-down predator-prey direction (Lynx → Hare).

**Figure 5.**
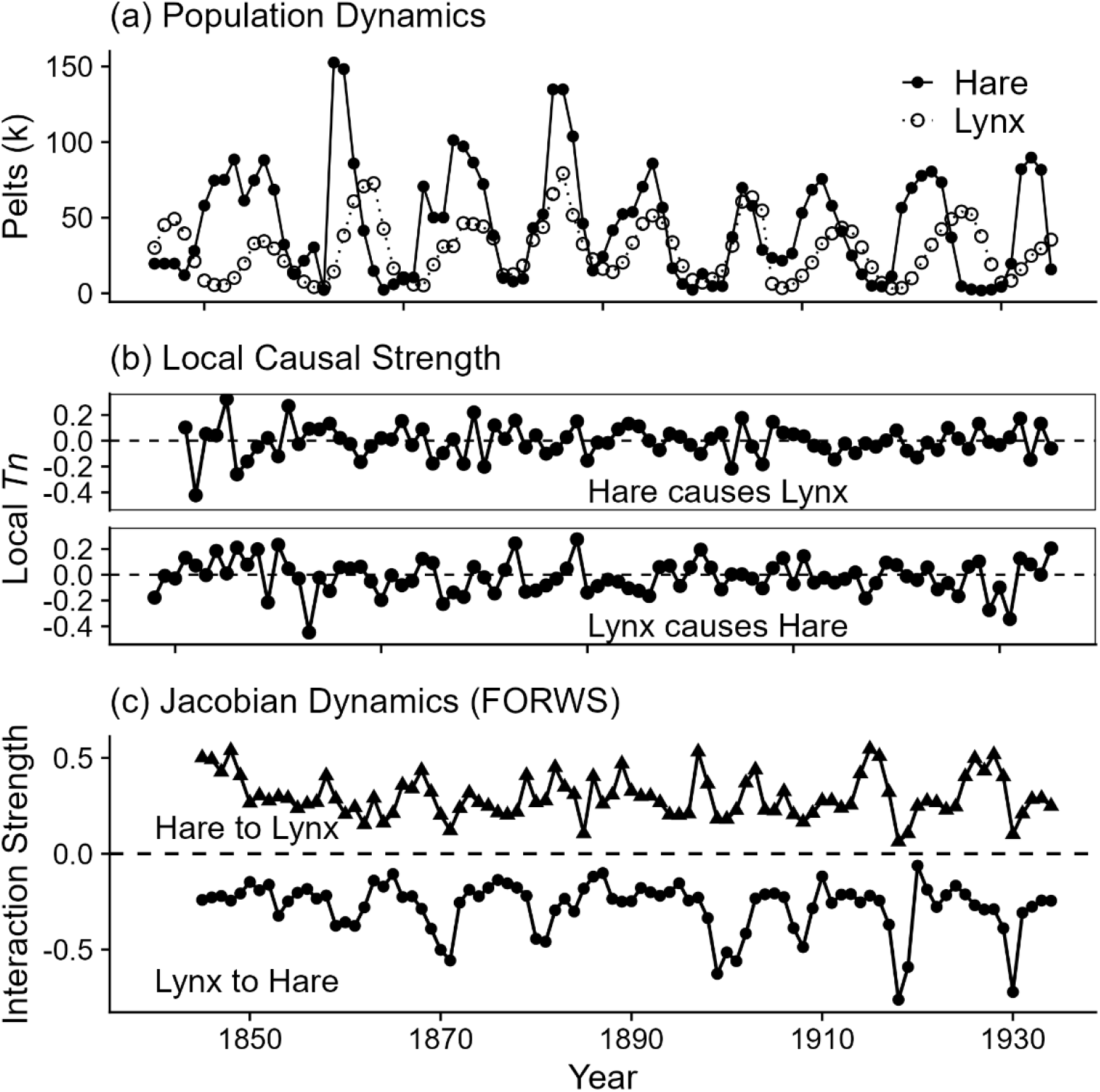
Multi-decadal population cycles, time-resolved local causality, and dynamic Jacobian interactions between Canada lynx and snowshoe hare. (a) Empirical population time series. Historical annual pelt returns for snowshoe hare (solid line with filled circles) and Canada lynx (dotted line with open circles) from the Hudson’s Bay Company archives (1845–1935). (b) Time-resolved local causal strength (*T_n_*). Rolling-window F-CODEC estimates (21-year moving window) for Hare → Lynx (top panel) and Lynx → Hare (bottom panel). Horizontal dashed lines mark the baseline *T_n_* = 0. Both directions exhibit episodic bursts of predictive coupling synchronized with cyclic population peaks and collapses. (c) State-dependent Jacobian interaction dynamics estimated by FORWS. Time-varying interaction strengths representing the per-capita effect of prey on predator recruitment (∂*Lynx_t_*_+1_/∂*Hare_t_*, top line with filled triangles) and predator impact on prey mortality (∂*Hare_t_*_+1_/∂*Lynx_t_*, bottom line with filled circles). The horizontal dashed line denotes zero interaction. FORWS consistently identifies positive trophic benefits for lynx and persistent negative predation pressure on hares across all cyclic regimes without requiring parametric functional response assumptions.

Time-resolved rolling-window analysis (21-year window) revealed that directional causal strengths fluctuated over time, exhibiting transient surges corresponding to major cyclic amplitude expansions around 1850–1870 and 1880–1900 (Fig. 5b). State-dependent Jacobian reconstruction via FORWS estimated time-varying interaction strengths throughout the 91-year series (Fig. 5c). The per-capita effect of prey on predator recruitment (∂Lynx_t+1_/∂Hare_t_) remained positive throughout the cycles (ranging from +0.06 to +0.55), whereas the predation impact on prey mortality (∂Hare_t+1_/∂Lynx_t_) remained persistently negative (ranging from –0.06 to –0.77). Predation pressure intensified sharply during prey population collapses (e.g., around 1870, 1900, and 1918) (Fig. 5a, c). Though both directional causal strengths and interaction strengths (Jacobian) fluctuated over time, the dynamics did not synchronize.

### Real-Time Tracking of Microclimate-Acoustic Causality and Collective Behavior in Honeybee Hives

In the in-situ IoT streaming monitoring of an active honeybee (*Apis mellifera*) colony, multivariable rolling-window F-CODEC tracked time-varying causal forcing between deseasonalized microclimate perturbations and hive acoustic responses across May and June 2026 (Fig. 6). The ambient temperature perturbation (T̃*emp_t_*_−1_) showed episodic causal coupling on hive sound (S̃*ound_t_*), with *T_n_* peaking above the local analytical threshold (*T*_crit_Win_ ≈ 0.056) in late May (reaching *T_n_* = 0.106) and immediately preceding an aborted swarming event on June 4 (*T_n_* = 0.104) (Fig. 6a). This benchmark event corresponded to an aborted swarming attempt, which is characterized by transient collective agitation and intensive flight activity that ultimately did not culminate in colony departure, serving as an in-situ biological ground truth for acute colony-level behavioral excitation. Hive volatile gas concentration (G̃*as_t_*_−1_) exhibited synchronized causal bursts (*T_n_* peaking at 0.105 and 0.091) during colony metabolic transitions (Fig. 6b).

**Figure 6.**
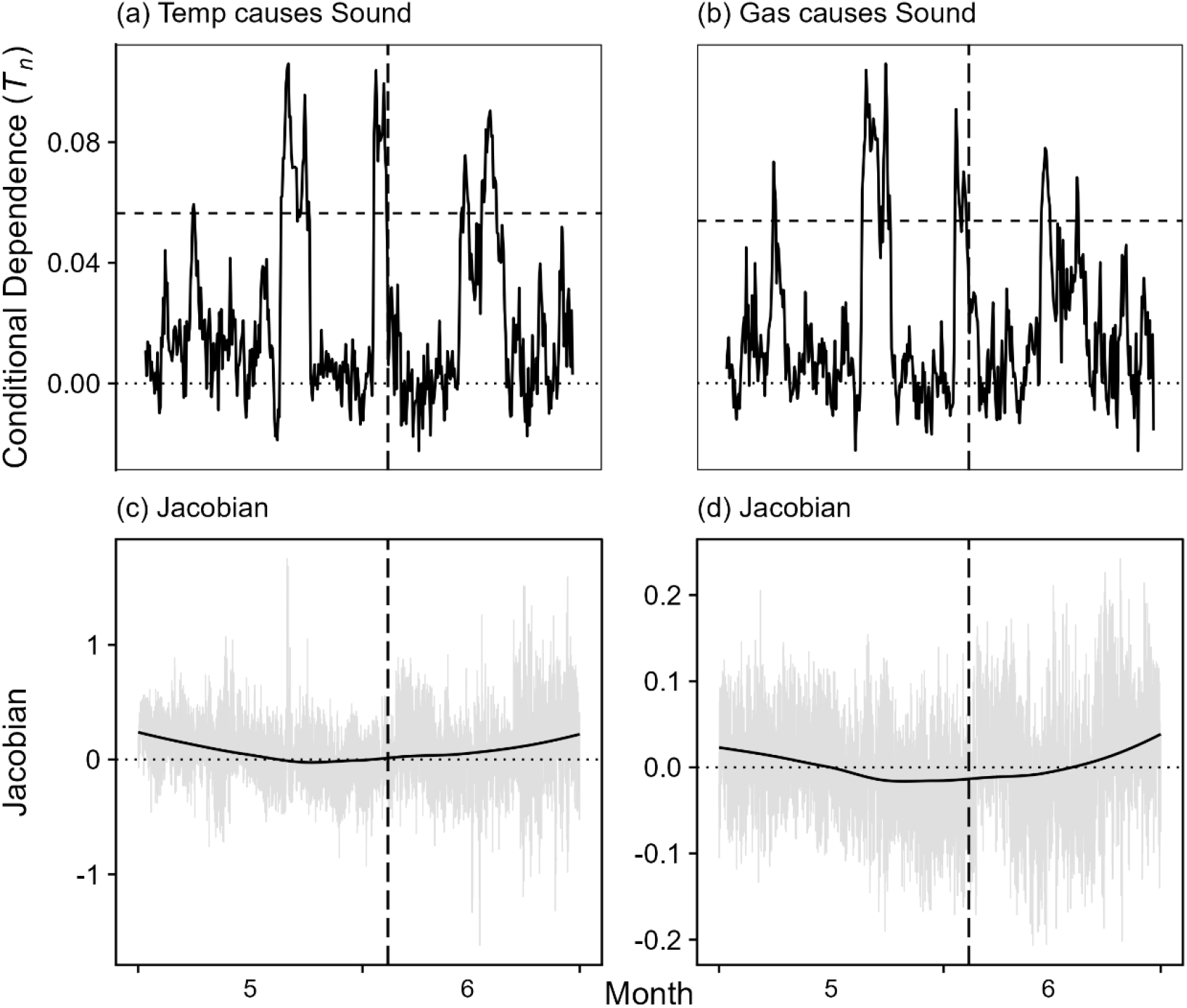
High-frequency in-situ IoT monitoring of an *Apis mellifera* hive across May and June 2026. The vertical dashed line marks an aborted swarming event recorded on June 4, 2026 (12:00 UTC). (a) Time-varying causal influence of temperature on acoustic response (*T_n_*). Rolling-window F-CODEC estimates (48-hour window, 3-hour step) evaluating the forward predictive contribution of prior temperature fluctuations (T̃*emp_t_*_−1_) on collective hive sound (S̃*ound_t_*). The horizontal dashed line denotes the local analytical 95% significance threshold (*T*_crit_Win_ ≈ 0.056); the horizontal dotted line indicates zero dependence (*T_n_* = 0). Temperature forcing exhibits episodic causal surges exceeding the critical threshold during late May and immediately preceding the June 4 event. (b) Time-varying causal influence of hive gas concentration on acoustic response (*T_n_*). Rolling-window F-CODEC evaluating prior volatile gas fluctuations (G̃*as_t_*_−1_) on hive sound. Gas concentration shows strong, synchronized bursts of causal coupling that peak sharply during specific colony metabolic and ventilation transitions. (c) Instantaneous and seasonal temperature Jacobian dynamics. Pointwise Jacobian interaction coefficients (*δS̃ound_t_*/*δ*T̃*emp_t_*_−1_) estimated by FORWS (grey lines) and long-term trend estimated by LOESS smoothing (solid black line). The horizontal dotted line represents zero interaction. (d) Instantaneous and seasonal gas Jacobian dynamics. Pointwise Jacobian interaction coefficients (*δ*S̃*ound_t_*/*δ*G̃*as_t_*_−1_) estimated by FORWS (grey lines) and LOESS seasonal trend (solid black line). Both Jacobians reveal state-dependent shifts from early-season active thermoregulation and fanning toward altered collective responsiveness following the June 4 swarming event.

Pointwise Jacobian sensitivities estimated by FORWS captured state-dependent behavioral shifts across the two-month period (Fig. 6c, d). Instantaneous temperature sensitivities (*δ*S̃*ound_t_*/*δ*T̃*emp_t_*_−1_) fluctuated around baseline zero in mid-May and shifted toward positive coupling during early summer warming (Fig. 6c). Instantaneous gas sensitivities (*δ*S̃*ound_t_*/*δ*G̃*as_t_*_−1_) exhibited negative interaction coefficients during late May (LOESS seasonal minimum of –0.0158 occurring at 12:58 UTC on May 26, 2026), consistent with previously reported collective ventilation and fanning behaviors (Fig. 6d).

### Discussion

Unraveling directional causality in complex dynamical systems has long been hindered by the dual challenges of generalized synchronization and prohibitive computational demands. Conventional state-space reconstruction methods, notably Convergent Cross Mapping (CCM), rely on backward-looking shadow manifold projections that collapse into mutually diffeomorphic geometries under strong dynamical coupling, inevitably producing symmetric false-positive detections. In contrast, F-CODEC reformulates causal discovery as a strictly forward-looking conditional dependence problem. By nonparametrically conditioning on the target variable’s intrinsic delay-coordinate history (**Z**_cond_), the target’s autonomous attractor manifold is projected out. This operation systematically absorbs the shared geometric structures induced by generalized synchronization, preventing spurious reverse causal inferences without requiring time-consuming surrogate data generation. Furthermore, the formulation of the rank-space manifold contraction factor (*γ*_Z_) and the effective degrees of freedom (*N*_eff_) provides a computationally instant, analytically motivated significance threshold directly from raw time series, transforming causal detection into an instantaneous, computationally efficient procedure.

A key architectural advance of F-CODEC lies in resolving the severe vulnerability of classical nonparametric metrics to high-dimensional observational noise. Standard conditional dependence measures, such as raw CODEC, rely on unweighted Euclidean 1-nearest-neighbor matching, which succumbs to distance concentration and topological distortion when the conditioning space contains non-informative or noisy variables. As demonstrated in our benchmark simulations, raw CODEC suffered a catastrophic loss of statistical power once uninformative noise dimensions were introduced. F-CODEC overcomes this limitation by leveraging Generalized Random Forests (GRF) to extract adaptive similarity weights. Because tree splits recursively prioritize coordinates that maximize conditional variance reduction, forest kernels naturally shrink along informative manifold dimensions while filtering out irrelevant noise coordinates. Moreover, replacing single-point nearest-neighbor matching with a normalized weighted expectation across multiple local neighbors provides essential rank smoothing, ensuring robust noise tolerance and high sensitivity even across sparse or non-uniformly distributed attractors.

Beyond pairwise interactions, reconstructing complex network architectures requires disambiguating direct physical drivers from indirect transitive cascades and common-cause confounding. In our multi-species benchmark motifs, F-CODEC proved highly effective in network pruning. In the trophic cascade chain (*X* → *Y* → *Z*), conditioning on the intermediate mediator *Y* neutralized the apparent transitive dependence between *X* and *Z*, reducing the metric to baseline zero. Similarly, in the common-cause fan-out motif (*Y* ← *X* → *Z*), conditioning on the shared parent *X* successfully eliminated spurious co-dependencies between *Y* and *Z*. These results confirm that multi-variable F-CODEC serves as an exact nonparametric counterpart to partial correlation or conditional transfer entropy, establishing a rigorous foundation for scaling automated causal graph discovery across high-dimensional interconnected systems without assuming linear Gaussian dynamics.

Application to the multi-decadal Canada lynx and snowshoe hare cycles highlights F-CODEC’s capability to resolve non-stationary, delayed ecological interactions. The bidirectional forward lag scan unambiguously identified an asymmetric top-down predatory forcing from lynx to hare peaking at a characteristic 2-year lag, whereas direct bottom-up forcing was not statistically detectable at the temporal resolution of annual pelt indices. This asymmetry likely reflects the fact that lynx population inertia and delayed density dependence mask short-term prey-to-predator recruitment signals in historical harvest records. Furthermore, integrating rolling-window F-CODEC with Forest-weighted S-maps (FORWS) provided complementary insights into ecosystem stability. While FORWS tracked persistent, state-dependent negative predation pressure (∂Hare_t+1_/∂Lynx_t_) that intensified during cyclic population crashes, rolling-window *T_n_*(*t*) quantified the episodic surges in directional predictive coupling. Crucially, our findings reveal that local causal strength (*T_n_*) and physical interaction sensitivity (Jacobian elements) are decoupled in phase, demonstrating that strong physical interactions do not always translate to high instantaneous predictability during non-equilibrium regime shifts.

The continuous monitoring of an active honeybee colony demonstrates the operational viability of F-CODEC for high-frequency IoT streaming analytics. By implementing a Double Machine Learning (DML) orthogonalization scheme with harmonic diurnal coordinates, we eliminated ubiquitous 24-hour circadian confounding, allowing multivariable rolling F-CODEC to detect genuine, short-term physiological perturbations. The framework successfully captured transient causal surges in temperature and gas forcing that preceded colony-level behavioral events, including an aborted swarming attempt and collective ventilation responses. Because F-CODEC bypasses computationally expensive surrogate simulations, it can be embedded directly into edge-computing microcontrollers and agricultural cyber-physical systems. In commercial smart greenhouses and precision livestock facilities, this enables real-time, closed-loop model predictive control that adapts dynamically to non-linear biological feedbacks rather than relying on static environmental thresholds.

Despite its robust performance, several avenues remain for extending the F-CODEC framework. First, while forest kernels handle moderate-to-high conditioning dimensions effectively, scaling to thousands of simultaneous variables—such as whole-transcriptome or global climate arrays—will benefit from integrating multi-view attractor distance ensembles and GPU-accelerated forest implementations. Second, our current formulation assumes discrete, regularly sampled time series; incorporating continuous-time neural ordinary differential equations or irregular kernel smoothing could extend F-CODEC to sparsely sampled longitudinal records. Finally, pairing F-CODEC with deep decision ensembles (such as boosted gradient trees) holds promise for identifying higher-order, multi-variable synergistic interactions. In summary, F-CODEC establishes a practical, scalable, and surrogate-free paradigm for causal discovery, bridging the gap between empirical dynamic modeling, machine learning, and real-time dynamical systems control. Furthermore, as with all conditioning-based causal discovery frameworks, F-CODEC operates under the standard assumption of causal sufficiency; while it rigorously prunes indirect pathways and spurious co-dependencies mediated by variables explicitly included in **Z**_cond_, latent or unobserved environmental confounders that force both systems externally remain an inherent challenge that warrants future integration with latent-variable modeling.

## Methods

### Mathematical Formulation of F-CODEC

#### Original Conditional Dependence Coefficient (CODEC)

Let *Y* be a one-dimensional continuous response variable, *X* be a *d*-dimensional candidate causal predictor vector, and *Z* be a *k*-dimensional vector of conditioning covariates observed across *n* temporal samples. The original Azadkia–Chatterjee conditional dependence coefficient quantifies the degree to which *Y* depends on *X* given *Z* without assuming parametric models or distributional shapes.

In rank space, let *R_i_* denote the rank of *Y_i_* among all *n* observations, taking integer values from 1 to *n*. For each sample point *i*, let *N_Z_*(*i*) and *N_XZ_*(*i*) represent the indices of the Euclidean 1-nearest neighbors of *Z_i_* in *Z*-space and of (*X_i_*, *Z_i_*) in the combined (*X*, *Z*)-space, respectively, excluding self-matches (*j* ≠ *i*):

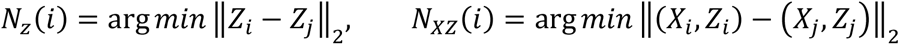

The empirical sample statistic *T_n_*(*Y*, *X* | *Z*) measures the relative preservation of target ranks across these topological nearest-neighbor pairs:

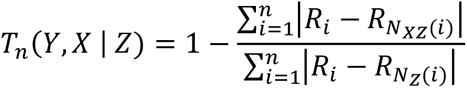

Under the null hypothesis where *Y* is conditionally independent of *X* given *Z*, *T_n_* converges to 0 as *n* approaches infinity. Conversely, when *Y* is a measurable, deterministic function of *X* given *Z*, *T_n_* approaches 1.

### Curse of Dimensionality in Nearest-Neighbor Matching

Standard CODEC relies on unweighted Euclidean nearest-neighbor matching. In nonlinear dynamical systems, rigorous state reconstruction requires multi-lag delay-coordinate embeddings or multivariable covariates, expanding the dimension of *Z* to *E*. As the embedding dimension *E* increases, the contrast between the distance to the nearest neighbor and the distance to the farthest neighbor diminishes rapidly, a phenomenon known as distance concentration. Furthermore, when observational noise or uninformative lag dimensions are present, unweighted Euclidean distances treat all coordinates (dimensions) equally, distorting the local neighborhood structure in *Z*-space and inflating false-positive causal detections, particularly when the samples are unevenly distributed in a state-space ^20^.

### Adaptive Forest Kernels via Generalized Random Forests

To overcome the distortion of Euclidean metrics in high-dimensional state spaces, F-CODEC replaces rigid 1-nearest-neighbor matching with adaptive similarity weights generated by Generalized Random Forests (GRF). Nonparametric regression forests are trained to predict the target *Y*: an autonomous forest trained solely on the conditioning set *Z*, and a joint forest trained on the combined set (*X*, *Z*).

For an ensemble of *B* decision trees, the raw similarity weight *α_ij_* between sample *i* and sample *j* is defined by the frequency with which they fall into the same terminal partition leaf *L_b_* across all trees (excluding self-matches):

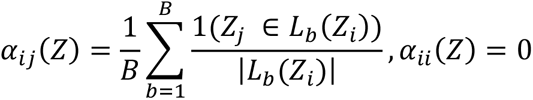

Because tree splits recursively maximize target variance reduction along the most informative coordinates, forest weights automatically shrink along dynamically relevant manifold dimensions while filtering out uninformative noise dimensions. To enforce topological sharpness and eliminate diffuse long-range weights, raw forest weights are thresholded to retain only the top *κ*-proportion (default *κ* = 0.20 or median cutoff) of local neighbors for each observation:

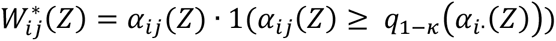

The filtered weights are then row-normalized to sum to unity:

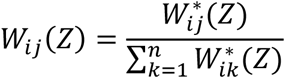

### F-CODEC Statistic Formulation

While the original CODEC calculates rank dispersion relying strictly on a single 1-nearest neighbor (*N_Z_*(*i*)), which makes it highly sensitive to local observational noise, F-CODEC estimates the expected local variation of the target *Y* as a weighted average across multiple neighboring samples determined by the adaptive forest weights.

Let *D_Y_*(*i*, *j*) denote the pairwise distance in the target space, defined as either the Euclidean distance |*Y_i_* − *Y_j_*| or the rank distance |*R_i_* − *R_j_*|. The expected local variation *D̂_i_* of the target *Y* around observation *i*, conditioned on *Z* alone and on (*X*, *Z*) jointly, is formulated by kernel smoothing over all candidate neighbors:

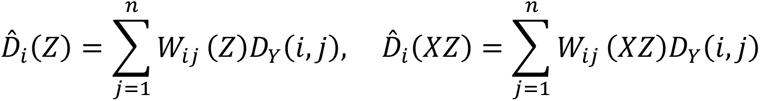

Here, because the filtered weights *W_ij_* are normalized such that ∑*_j_ W_ij_* = 1, *D̂_i_* represents the weighted conditional expectation of target differences within the adaptive topological neighborhood of sample *i*. The Forest-weighted Conditional Dependence Coefficient (F-CODEC) is then defined as the aggregate relative reduction in this expected local dispersion:

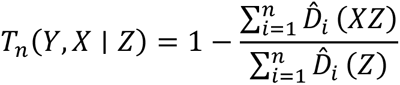

For dynamical causal inference, F-CODEC configures all variables strictly forward in time. Specifically, it evaluates the unique predictive information that a time-lagged causal candidate contributes to forecasting the future state of the target variable, beyond what is already captured by the target’s own historical delay-coordinate trajectory. By nonparametrically conditioning on this intrinsic attractor history, the target’s autonomous dynamical components are effectively projected out. This operation systematically neutralizes the shared geometric representations arising from generalized synchronization and network confounding, thereby isolating genuine directional causality without relying on computationally intensive surrogate time series.

To strictly prevent target leakage and adaptive overfitting, F-CODEC leverages the inherent honest splitting and out-of-bag (OOB) estimation protocol of generalized random forests (grf), wherein forest similarity weights for each sample are computed exclusively from trees where that observation was omitted during training, and self-referential weights are explicitly removed by setting diag(*W*) = 0.

### Analytical Significance Threshold for Time Series

Under the null hypothesis of conditional independence for i.i.d. observations, the sample statistic *T_n_* asymptotically follows a normal distribution with mean zero and standard error 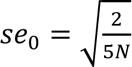. However, in deterministic dynamical systems, temporal autocorrelation and the deterministic structure of the conditioning attractor manifold reduce the effective degrees of freedom and contract the residual variance of the target variable. To establish an instant hypothesis test without surrogate data, F-CODEC incorporates a two-fold correction. First, the effective sample size *N*_eff_ is estimated using the lag-1 autocorrelation (*ρ*_1_) of the target time series:

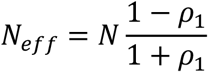

Second, the degree of manifold contraction is quantified in rank space. Let *R_Y_* be the rank vector of Y, and *R_Z_* be the rank-transformed coordinates of **Z**_cond_. We estimate the multiple coefficient of determination (*R*^2^_rank_) by linear projection of *R_Y_* onto *R_Z_*. The residual variance contraction factor is defined as: *γ*_Z_ = 1 – *R*^2^_rank_, with a lower threshold imposed at *γ*_Z_ ≥ 0.05 to safeguard against asymptotic variance collapse in strictly deterministic regimes.

The analytical, autocorrelation-corrected 95% significance threshold is formulated as:

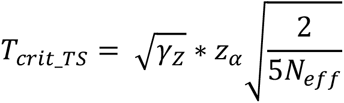

where z*_α_* = 1.645 for nominal *α* = 0.05. A directional causal link is deemed statistically significant when *T_n_* > *T*_crit_TS_. Because *T_n_* measures the relative variance reduction contributed by *X*, the causal hypothesis test is formulated strictly as a one-sided upper-tail test (*H*_0_: *T_n_* ≤ 0 versus *H*_1_: *T_n_* > 0). In deterministic dynamics, conditioning on **Z**_cond_ strongly contracts the denominator *D_i_*(*Z*), whereas appending an uninformative candidate *X* slightly dilutes nearest-neighbor locality in finite samples, shifting the empirical null distribution toward negative values (Supplementary Fig. S1). The resulting empirical test size is strictly bounded by nominal *α*, ensuring robust protection against false discoveries without sacrificing statistical power in the presence of true coupling (Supplementary Fig. S2).

### Dynamic Interaction Sensitivity via Forest-Weighted S-Maps (FORWS)

To track time-varying Jacobian interaction strengths (partial derivatives) across state space, we integrate the adaptive forest similarity weights into a regularized local linear regression framework (Forest-weighted S-map; FORWS).

Let *X*_full_ be the concatenated state matrix including an intercept column, and *Y*_full_ be the one-step-ahead target vector. For each observation point *k*, the local interaction coefficient vector *c_k_* is estimated by minimizing the weighted penalized objective:

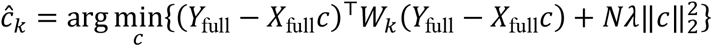

where *W_k_* = diag(*W_k_*_1_, …, *W_kN_*) is the diagonal weight matrix derived from a regression forest trained on the state coordinates, and *λ* is the Tikhonov regularization parameter (default *λ* = 10^−4^). The analytical solution is obtained via generalized ridge inversion:

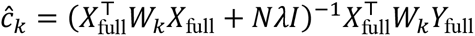

The resulting time series of model coefficients *ĉ_k_* provides the instantaneous Jacobian sensitivities (e.g., *δY_t_*_+1_/*δX_t_*), capturing state-dependent interaction strengths without parametric model specification.

### Synthetic Benchmark on Coupled Chaotic Maps Coupled Map Simulation Protocol

To evaluate the directional causal inference capability and robustness of F-CODEC against generalized synchronization and uninformative dimensions, we constructed a unidirectionally coupled chaotic logistic map system:

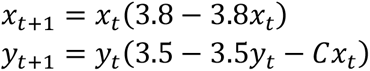

where *C* ∈ [0.0, 0.7] denotes the unidirectional coupling strength from *x* to *y*. The system was iterated for 400 time steps, discarding the initial 100 burn-in iterations to remove transient dynamics and retain a sample length of *N* = 300. The state variables were bounded within [0.0001, 0.9999] to ensure numerical stability. Observational noise was introduced by adding zero-mean Gaussian perturbations with a standard deviation equal to 5% of the standard deviation of each raw time series (*σ*_noise_ = 0.05 × SD). To simulate high-dimensional environmental confounding and test susceptibility to the curse of dimensionality, five independent standard normal random noise variables were generated and appended as dummy covariate dimensions (noise_lag_1_ to noise_lag_5_).

### Quantitative Metric Dynamics across Coupling Strengths

We examined the continuous response of the F-CODEC statistic *T_n_* and its empirical confidence intervals across varying coupling strengths (*C* = 0.0 to 0.5 in increments of 0.1) over 15 independent realization trials.

To assess causality direction skill, each simulation trial was evaluated for both the true causal link (*X* → *Y*) and the non-causal reverse link (*Y* → *X*). In the true causal direction, the target state was designated as *y_t_*_+1_, the causal candidate predictor as *x_t_*, and the conditioning set **Z**_cond_ as the target’s intrinsic history *y_t_* concatenated with the five dummy noise variables. Conversely, for the spurious reverse direction, the target state was configured as *x_t_*_+1_, the candidate predictor as *y_t_*, and the conditioning set **Z**_cond_ as *x_t_* along with the five noise variables.

F-CODEC was computed using *B* = 1000 regression trees, a minimum leaf node size of 5, and a top 20% local weight threshold (*κ* = 0.20). Mean *T_n_* values and empirical 95% percentile confidence intervals (2.5th and 97.5th percentiles) were aggregated across trials. The asymptotic analytical 95% significance threshold under the null hypothesis was calculated as 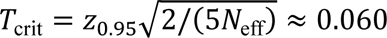 for effective sample size *N*_eff_ = 299.

### Comparative Benchmark Across Causal Inference Methods

To evaluate causal detection performance, we systematically evaluated causal detection rates (statistical power and false-positive control at nominal *α* = 0.05) across coupling strengths ranging from *C* = 0.0 to 0.7 using multiple independent realization trials (50 trials per condition for F-CODEC and Raw CODEC; 200 trials for FORCC and CCM). This comparison evaluated six methodological configurations: (i) F-CODEC without noise, conditioning on the target’s intrinsic dynamics **Z**_cond_ = [*y_t_*, *y_t_*_–1_] with optimal embedding dimension *E* = 2; (ii) Raw CODEC without noise, faithfully employing classical Euclidean 1-nearest-neighbor matching across the identical 2-dimensional conditioning set; (iii) F-CODEC with 5 noise dimensions, conditioning on the expanded vector **Z**_cond_ =[*y_t_*, *y_t_*_–1_, noise_lag_1_,…, noise_lag_5_]; (iv) Raw CODEC with 5 noise dimensions, searching Euclidean nearest neighbors across the 7-dimensional space; (v) FORCC (Chatterjee), evaluating forward causality using regression-forest-weighted Chatterjee rank correlation (Kishi unpublished); and (vi) conventional Convergent Cross Mapping (CCM), performing backward state-space cross-mapping based on Takens’ delay embedding and simplex projection. For each trial, causal detection was registered as a binary success when the estimated test statistic exceeded the theoretical 95% threshold *T*_crit_ for F-CODEC and CODEC, or the 95% surrogate significance threshold for CCM.

### Three species Dynamical System

To evaluate the capability of F-CODEC in distinguishing direct causal links from indirect transitive cascades and common-cause spurious correlations, we constructed two canonical three-species interaction motifs (*N* = 300 time steps after discarding 100 burn-in steps; 50 independent realization trials per motif). One is a multi-trophic cascade chain (*X* → *Y* → *Z*). Species *X* and *Y* adversely affect *Y*, and *Z*, respectively.

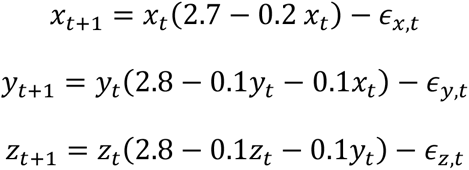

The other is a fanout relationship (*Y* ← *X* → *Z*). Species *X* exerts a negative effect on both *Y* and *Z*.

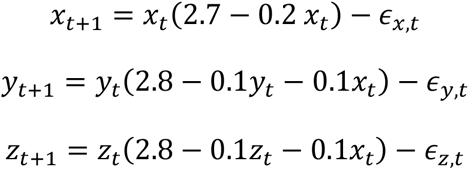

In both systems, independent stochastic process noise perturbations, *∊*_.,t_ were sampled from a normal distribution with a mean value 0 and standard deviation 0.05 and added at each time step.

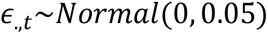

Initial conditions were assigned as *x*_1_ = 8.0, *y*_1_ = 9.0 and *z*_1_ = 3.5.

During numerical simulation, state values falling below zero were bounded to a lower floor of 0.001 to prevent non-physical negative population densities. For each motif, we systematically evaluated all six pairwise directional permutations among *X*, *Y*, and *Z* using an embedding dimension of *E* = 2. All predictor and conditioning matrices were standardized to zero mean and unit variance prior to forest construction. The F-CODEC statistic *T_n_* was calculated using *B* = 1000 regression trees with a top 20% local neighbor filter (*κ* = 0.20). Significance for each direction was tested against the rank-space autocorrelation-corrected threshold *T*_crit_TS_ calculated with effective degrees of freedom *N*_eff_ = 298 at nominal *α* = 0.05.

### Empirical Data Acquisition and Preprocessing

To evaluate the performance of F-CODEC on real-world non-equilibrium ecological dynamics, we utilized the classic multi-decadal time series of Canada lynx (*Lynx canadensis*) and snowshoe hare (*Lepus americanus*) pelt records from the Hudson’s Bay Company archives spanning 1845 to 1935 (*N* = 91 annual observations). The digitized dataset was downloaded from a public “Github” repository (see Data Availability). Prior to state-space reconstruction and causal calculations, both time series were standardized to zero mean and unit variance.

### Optimal Embedding Dimension and Bidirectional Forward Lag Scanning

We first determined the optimal embedding dimension *E*\* for each species autonomously by scanning *E* ∈ [1, 8] using F-CODEC. Specifically, for a candidate dimension *e*, the future state *Y_t_*_+1_ was predicted from the immediate past *Y_t_*, nonparametrically conditioned on the remaining historical embedding coordinates **Z**_cond_ = [*Y_t_*_-1_, …, *Y_t_*_–*e*+1_]. The optimal dimension *E*\* was selected where the unconditional predictive gain *T_n_* reached its global maximum (yielding *E*\* = 2 for lynx and *E*\* = 2 for hare).

Using these embedding dimensions, we performed bidirectional forward lag scans across time delays L∈ [0, 8] years. To evaluate the bottom-up hypothesis (Hare → Lynx), the target was set to *Lynx_t_*_+1_ conditioned on its own intrinsic trajectory **Z**_cond_ = [*Lynx_t_*, …, *Lynx_t_*_–*E*\*+1_], while testing candidate predictor vectors **X***_t_*_–*L*_ = [*Hare_t_*_–*L*_, …, *Hare_t_*_–*L*–*E*\*+1_]. Conversely, for the top-down predator-prey hypothesis (Lynx → Hare), the target was set to *Hare_t_*_+1_ conditioned on [*Hare_t_*, …, *Hare_t_*_–*E*\*+1_], driven by lagged vectors **X***_t_*_–*L*_ = [*Lynx_t–L_*, …, *Lynx_t_*_–*L–E*\*+1_]. Directional significance for each lag was assessed against the rank-space autocorrelation-corrected threshold *T*_crit_TS_ at nominal *α* = 0.05. Because scanning across multiple candidate time delays (*L* ∈ [0, 8] years) involves simultaneous hypothesis evaluations, we addressed potential family-wise false-positive inflation by verifying candidate peaks against a conservative Bonferroni-corrected threshold (*α*_Bonf_ = 0.05/9 ≈ 0.0055, corresponding to critical threshold *T*_crit_Bonf_), ensuring that identified causal delays reflect genuine dynamical coupling rather than multiple-testing artifacts.

To investigate how causal forcing and ecological interaction strengths varied over the 91-year period, we applied rolling-window F-CODEC and the Forest-weighted S-map (FORWS). Time-resolved conditional dependence *T_n_*(*t*) was calculated across a moving window of 21 years (half-width of 10 years). Within each temporal window, local F-CODEC was evaluated for the optimal lags identified in the global lag scan, tracking the temporal fluctuations of directional predictability. To quantify dynamic interspecific interaction strengths, we constructed a two-variable block model predicting one-step-ahead standardized densities [*Hare_t_*_+1_, *Lynx_t_*_+1_] from current states [*Hare_t_*, *Lynx_t_*]. A multi-output regression forest (*B* = 1000 trees) was trained on the predictor matrix to extract state-dependent forest similarity weights *W_ij_* for each historical observation *i*. Using these local weights, we solved regularized ridge regressions (penalty parameter *λ* = 10^−4^) to compute the time-varying Jacobian matrix elements: ∂*Hare_t_*_+1_/∂*Lynx_t_* (predation impact) and ∂*Lynx_t_*_+1_/∂*Hare_t_* (trophic benefit).

### IoT Environmental and Acoustic Monitoring Protocol

To validate the utility of F-CODEC in tracking real-time, non-stationary physiological responses in complex biological systems, we deployed an autonomous environmental and acoustic monitoring device inside an active European honeybee (*Apis mellifera*) hive. High-frequency time series spanning May 1 to June 30, 2026, were collected, recording internal nest temperature (°C), volatile gas concentration, and continuous hive audio recordings. Raw acoustic audio files were pre-processed into 100-dimensional spectral feature vectors per time window. To extract the primary collective behavioral state from acoustic features while filtering high-frequency recording anomalies, Uniform Manifold Approximation and Projection (UMAP; *n_neighbors_* = 15, *min_dist_* = 0.1) was applied to map the acoustic spectra into a low-dimensional manifold. The primary inverted axis (Sound U1) was defined as the collective acoustic activity index. During the two-month deployment, a naturally occurring aborted swarming event was visually recorded at 12:00 UTC on June 4, 2026, providing an empirical benchmark of sudden collective behavioral reorganization.

### Diurnal Cycle Deseasonalization via Double Machine Learning

Circadian fluctuations in ambient solar irradiance drive strong diurnal rhythms across hive temperature, gas exchange, and worker activity, acting as ubiquitous periodic confounders. To eliminate spurious causal associations driven purely by shared 24-hour cycles, we implemented a Double Machine Learning (DML) orthogonalization scheme using nonparametric regression forests. Harmonic diurnal basis coordinates were constructed as *Z*_time_ = [sin(2*πt*/24), cos(2*πt*/24)], where *t* denotes the fractional hour of the day. Separate regression forests (*B* = 300 trees) were trained to predict each raw variable from *Z*_time_:

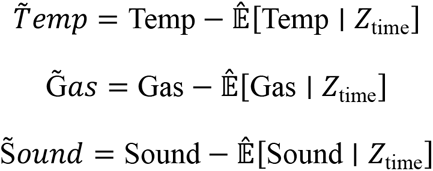

These orthogonalized residuals isolate non-circadian physiological perturbations, reflecting true endogenous colony responses and exogenous weather events.

### Rolling-Window Multivariable F-CODEC Dynamics

To track time-varying causal forcing on colony sound dynamics, we applied a rolling-window multivariable F-CODEC across the deseasonalized streaming time series. A moving temporal window of 48 hours (nominally 384 observations at 7.5-minute intervals, averaging an effective sample size of *N_win_* = 340 after quality control and removal of non-concurrent records) was shifted in 3-hour steps (24 consecutive observations) across the two-month monitoring period. Within each analysis window, the target variable was defined as the current acoustic state (*S̃ound_t_*), and the putative causal drivers were set to the prior one-step lagged microclimate perturbations (*T̃emp_t_*_−1_, *G̃as_t_*_−1_). To project out the target’s intrinsic acoustic persistence and control for any residual diurnal baseline shifts, the conditioning set **Z**_cond_ incorporated the target lag-1 history alongside harmonic time coordinates: [*S̃ound_t_*_−1_, sin(2*πt*/24), cos(2*πt*/24)]. Multivariable F-CODEC was then evaluated for each window using *B* = 150 regression trees with median weight thresholding. The local significance threshold was evaluated analytically as 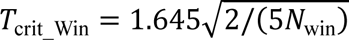, which corresponds to approximately 0.056 for *N_win_* = 340.

### Jacobian Dynamics via FORWS

To quantify the sign and magnitude of instantaneous sensitivities (*δS̃ound_t_*/*δ*T̃*emp_t_*_−1_ and *δ*S̃*ound_t_*/*δ*G̃*as_t_*_−1_), we applied the Forest-weighted S-map (FORWS). A regression forest (*B* = 500 trees) was trained on the joint lagged predictor matrix [Tẽmp*_t_*_−1_, G̃as*_t_*_−1_, Soũnd*_t_*_−1_] predicting S̃*ound_t_*. Dynamic similarity weights *W_ij_* were extracted for each observation point *i*, and regularized locally weighted linear models were fitted to estimate the continuous trajectory of Jacobian interaction coefficients across the two-month monitoring period. A localized LOESS smoother was applied to visualize the broad seasonal baseline shift of Jacobian sensitivities across early summer.

### Statistical analysis

All causal significance tests based on F-CODEC were formulated strictly as one-sided upper-tail hypothesis tests (*H*_0_: *T_n_* ≤ 0 versus *H*_1_: *T_n_* > 0) evaluated at a nominal significance level of *α* = 0.05. In time-series analyses, significance boundaries (*T*_crit_TS_) were corrected analytically using the effective sample size (*N*_eff_) based on lag-1 autocorrelation and the rank-space manifold contraction factor (γ_Z_). Multi-lag exploratory scans across delays L ∈ [0, 8] years were adjusted for multiple testing using conservative Bonferroni correction (*α*_Bonf_ = 0.05 / 9 ≈ 0.0055). In benchmark simulations, causal detection rates represent the proportion of trials where the sample test statistic exceeded the theoretical 95% critical threshold across independent realization trials (*n* = 50 or *n* = 200, specified per experiment). Pointwise interaction coefficients (Jacobians) and rolling-window conditional dependences are reported with local LOESS fits and empirical standard errors as detailed in the figure legends.

All statistical calculations, simulations, and data visualizations were executed in the R statistical computing environment (R version 4.3.0)^21^. Nonparametric regression forests and adaptive weight extraction were implemented using the grf package (version 2.3.0)^22^. Comparative benchmarks were executed using rEDM (version 0.7.5, URL: https://ha0ye.github.io/rEDM/articles/rEDM.html) for Convergent Cross Mapping and rUIC (version 0.9.15, URL: https://github.com/yutakaos/rUIC) for information-theoretic transfer entropy. Dimensionality reduction of acoustic spectra was conducted via uwot (version 0.1.16)^23^. Asynchronous parallel computing was orchestrated across multi-core architectures using the future and furrr packages.

### LLM Declarations

During the preparation of this manuscript, the author used Google Gemini to improve the English language, refine the readability of the text and R code. After using this tool, the author thoroughly reviewed and edited the content as needed and takes full responsibility for the final content of the publication.

## Data and code availability

The historical Canada lynx and snowshoe hare pelt records are accessible via the Living-Data-Tutorials repository (https://github.com/Living-Data-Tutorials/lynx_hare). The other data and custom R code supporting the findings of this study are available from the corresponding author upon reasonable request.

## Acknowledgments

I thank Dr. Takehiko Yamanaka for invaluable discussions and comments on the previous draft. A part of this research was conducted using the AI research supercomputer ‘SHIHOU’ at the National Agriculture and Food Research Organization (NARO). This study is partly supported by JSPS KAKENHI Grant Number JP23751015 and JP26K09087.

## Author contributions

S.K. is the sole author of this work. S.K. conceived the original idea, developed the theoretical framework, performed the data collection and analyses, and wrote the manuscript.

## Competing Interests

The author declares a competing financial interest in the form of a pending patent application in Japan related to the F-CODEC presented in this study, adding to FORWC, FORCC, and FORWS (Japanese Patent Application No. 2026–87255, titled “Prediction method, causal inference method, prediction apparatus, causal inference apparatus, and program”, by SK).

## Notes

https://github.com/Living-Data-Tutorials/lynx_hare

