## Supplementary figures S1-S3 for "Forward time-series causal inference based on forest weights and conditional dependence"

Shigeki Kishi*

### Overview of Supplementary Materials

This document provides supplementary analytical validations, robustness benchmarks, and computational efficiency profiles supporting the findings in the main text. There are 3 figures each with description.

- **Supplementary Note 1 (Figure S1):** Empirical null distribution calibration and Type I error rate verification across independent stochastic, autocorrelated, and chaotic processes.
- **Supplementary Note 2 (Figure S2):** Comprehensive two-dimensional parameter grid heatmap evaluating true-positive statistical power and false-positive resilience across varying dynamical coupling strengths and observational noise levels.
- **Supplementary Note 3 (Figure S3):** High-throughput computational runtime benchmarks comparing surrogate-free F-CODEC against 100-surrogate state-space (rEDM) and information-theoretic (rUIC) causal inference pipelines.

### Computational Environment and Software Reproducibility

All numerical simulations, empirical analyses, and benchmarking workflows were conducted in the R statistical computing environment R version 4.4.0 (R core team 2026). Asynchronous parallel execution was managed using future and furrr across available CPU cores. Major packages used in this supplementary analyses are grf, rEDM, rUIC, and XICOR. The "grf" package (Generalized Random Forests, version 2.3.0) (Tibshirani et al. 2026) was used for training honest regression trees, out-of-bag (OOB) sample splitting, and extracting adaptive similarity weights. The "rEDM" package (version 0.7.5, compiled from source via GitHub: ha0ye/rEDM) was used for Convergent Cross Mapping (CCM) and simplex projection ((Sugihara et al. 2012)). The "rUIC" version 0.9.15 (Unified Information Causality) (https://github.com/yutakaos/rUIC) and "macamts" version 0.2.5 (https://github.com/ong8181/macam/, DOI: 10.5281/zenodo.19244113) for multivariable state-space reconstruction and transfer-entropy-equivalent evaluations. The "XICOR" package was used for rank correlation baselines, especially for calculation of the Chatterjee correlation coefficient. Other packages were "uwot" for Uniform Manifold Approximation and Projection (UMAP), and "tidyverse" along with "cowplot" and "patchwork" for data synthesis and visualization.

**Supplementary Note 1: Empirical Null Calibration and Type I Error Control**

We examined if the analytical time-series threshold ($T_{\mathrm{crit\_TS}}$) suppresses false-positive when given an unrelated but noisy pair of time series without surrogate testing. We evaluated the empirical null distribution of $T_n$ across three canonical non-causal dynamical regimes ($N$ = 300, observations per trial, 500 independent realization trials per regime):

1. **Autoregressive AR(1) Process:** Pairs of mutually independent stochastic processes with strong temporal persistence,

$$x_t = 0.6x_{t-1} + \epsilon_{x,t}$$

$$y_t = 0.6y_{t-1} + \epsilon_{y,t}$$

where $\epsilon \sim \mathcal{N}(0,1)$. This benchmark assesses susceptibility to red-noise autocorrelation artifacts.

2. **Uncoupled Deterministic Chaos:** Two completely independent chaotic logistic maps,

$$x_{t+1} = 3.8x_t(1 - x_t)$$

$$y_{t+1} = 3.8y_t(1 - y_t)$$

initialized from independent random starting states ($x_1, y_1 \sim$ Uniform$(0.2,0.8)$). This evaluates metric stability in the presence of strong deterministic nonlinear attractor geometry.

3. **Gaussian White Noise:** Uncorrelated, independent standard normal sequences,

$$x_t \sim \mathcal{N}(0,1)$$

$$y_t \sim \mathcal{N}(0,1)$$

serving as the baseline independent and identically distributed benchmark.

For each realization, candidate causality was tested by configuring the target as $Y_{t+1}$ and the candidate driver as $X_t$, while nonparametrically conditioning on the target's own two-dimensional lag history $\mathbf{Z}_{\text{cond}} = [Y_t, Y_{t-1}]$ ($E^* = 2$). The analytical 95% threshold $T_{\text{crit\_TS}}$ was computed for each trial incorporating the lag−1 autocorrelation correction ($N_{\text{eff}}$) and the rank-space manifold contraction factor ($\gamma_Z$).

Across all 1,500 independent null simulations (500 per regime), the empirical distribution of $T_n$ remained strictly below the analytical significance boundary ($T_{\mathrm{crit\_TS}}$), yielding an empirical Type I error rate of 0.0% at nominal $\alpha = 0.05$ (Supplementary Fig. S1).

In Gaussian white noise and AR(1) processes, the empirical null distributions centered near zero (mean $T_n = -0.001$ and $-0.004$, respectively), with their upper tails comfortably bounded by $T_{\mathrm{crit\_TS}}$. In uncoupled chaos, the null distribution shifted decisively into the negative domain (mean $T_n = -0.147$, range [–0.21, –0.07]). This negative shift arises from the geometric properties of deterministic attractor conditioning: the intrinsic history $\mathbf{Z}_{\mathrm{cond}}$ accurately reconstructs the local manifold of $Y$, strongly contracting the baseline local dispersion $D_i(Z)$. Adding an uninformative, independent driver $X$ unavoidably inflates the local neighborhood diameter in finite samples—a manifestation of the finite-sample penalty for uninformative conditioning dimensions. Consequently, the joint local dispersion slightly exceeds the autonomous dispersion ($D_i(XZ) \geq D_i(Z)$), naturally driving the ratio ($D_i(XZ)/D_i(Z) > 1$) and yielding negative $T_n$ values under the null hypothesis.

Importantly, while an empirical Type I error rate of 0.0% indicates that the analytical threshold $T_{\mathrm{crit\_TS}}$ acts conservatively under strictly uncoupled regimes, this conservative behavior does not compromise statistical power (Type II error). As demonstrated in the two-dimensional parameter benchmark (Supplementary Fig. S2a) and main benchmark (Fig. 2c), the test achieves and maintains 100% true-positive sensitivity once genuine physical coupling is present ($C \geq 0.15$), even in the presence of 20% observational noise. The threshold thus acts as an ideal safeguard against generalized synchronization and autocorrelation-induced artifacts while preserving high sensitivity to true causal forcing.

**Figure S1**

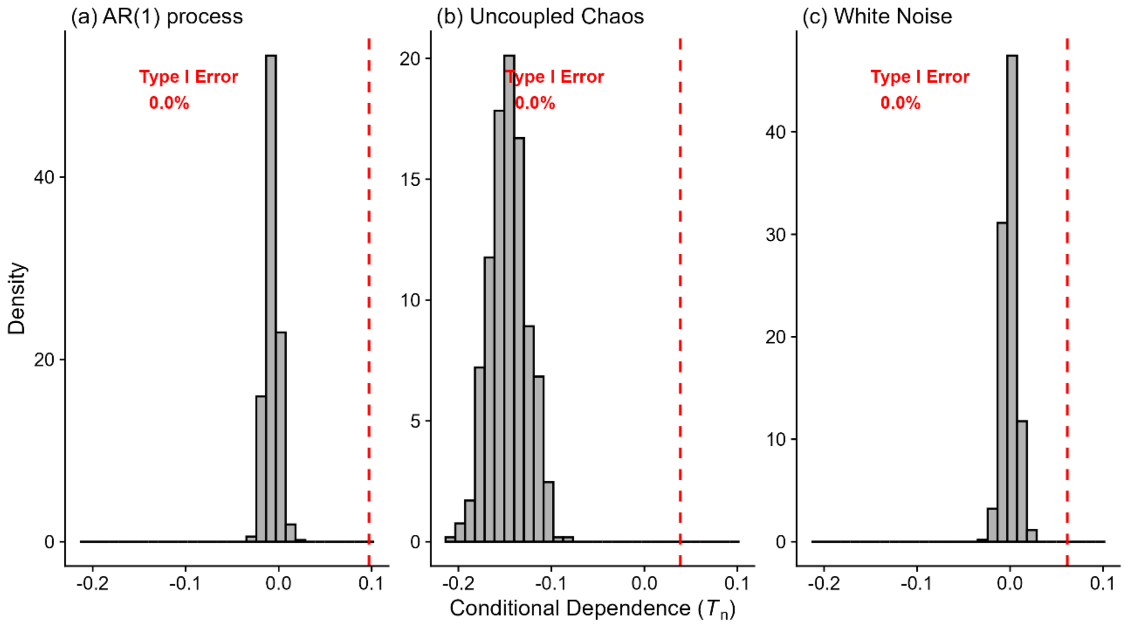


**Figure S1. Empirical null distributions of F-CODEC and Type I error calibration across distinct dynamical regimes.** Histograms of the sample conditional dependence coefficient ($T_n$) evaluated across 500 independent realization trials ($N = 300$ time steps) under the null hypothesis of no causal linkage. In all evaluations, target history coordinates $\mathbf{Z}_{\mathrm{cond}} = [Y_t, Y_{t-1}]$ were conditioned out. Vertical dashed red lines denote the mean analytical significance threshold ($T_{\mathrm{crit\_TS}}$) at nominal $\alpha = 0.05$.

**(a) Autoregressive AR(1) process.** Mutually independent stochastic series with lag-1 autocorrelation coefficient $\phi = 0.6$. The analytical threshold ($T_{\mathrm{crit\_TS}} \approx 0.098$) accounts for variance inflation due to autocorrelation, yielding an empirical Type I error rate of 0.0%.

**(b) Uncoupled deterministic chaos.** Two independent chaotic logistic maps ($C = 0.0$). Deterministic manifold conditioning shifts the empirical null distribution into negative values (mean $T_n \approx -0.147$), safely below the threshold ($T_{crit\_TS} \approx 0.041$), resulting in a 0.0% false-positive rate.

**(c) Gaussian white noise.** Independent and identically distributed standard normal noise series. The distribution centers at zero, bounded by the threshold ($T_{crit\_TS} \approx 0.063$), confirming 0.0% false-positive detections.

### Supplementary Note 2: Robustness Across Dynamical Coupling Strengths and Observational Noise Levels

To verify that F-CODEC operates robustly across diverse dynamical regimes rather than within narrow, hand-tuned parameter configurations, we evaluated true-positive detection sensitivity (statistical power) and false-positive resilience across a two-dimensional grid of dynamical coupling strengths and observational noise levels.

#### 2.1| Simulation Design and Parameter Space

Synthetic time series were generated using the unidirectionally coupled chaotic logistic map system ($X \rightarrow Y$, $N = 300$ observations per trial) across 55 parameter combinations:

- **Coupling Strengths ($C$):** Eleven levels ranging from $C = 0.00$ (uncoupled null regime) to $C = 0.50$ (strong coupling near complete dynamical synchronization) in increments of 0.05.
- **Observational Noise Levels ($\sigma_{\text{noise}}$):** Five noise-to-signal standard deviation ratios ($\sigma_{\text{noise}} \in \{0.01, 0.05, 0.10, 0.15, 0.20\}$), where independent Gaussian noise $\mathcal{N}\left(0, \sigma_{\text{noise}}^2 \cdot \text{Var}(X)\right)$ was added to both state variables.
- **Replication and Testing:** Twenty independent realization trials were executed for each parameter cell ($55 \times 20 = 1{,}100$ total simulations). For each realization, bidirectional causality was tested ($X \rightarrow Y$ and $Y \rightarrow X$) using an embedding dimension of $E^* = 2$, conditioning on the target's intrinsic delay coordinates $\mathbf{Z}_{\text{cond}} = [Y_t, Y_{t-1}]$. Detections were registered when the estimated $T_n$ exceeded the analytical time-series threshold $T_{\text{crit\_TS}}$ at nominal $\alpha = 0.05$.

#### 2.2| Grid Performance and Synchronization Resilience

The parameter landscape demonstrates sharp directional separation and robust noise tolerance across all tested regimes (Fig. S2).

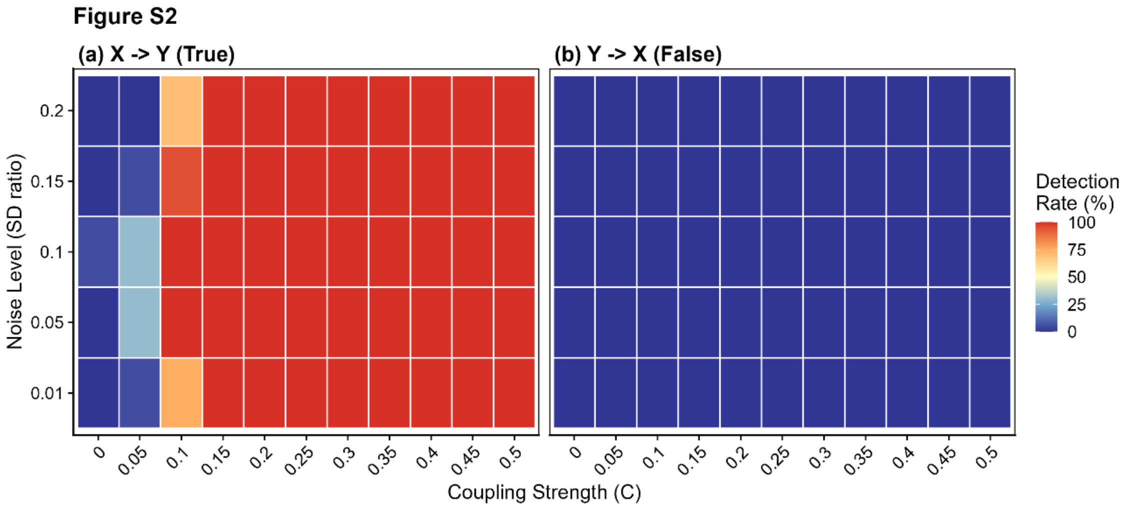


**Figure S2. Statistical power and false-positive rate grid across varying coupling strengths and observational noise levels.** Heatmaps of the causal detection rate (%) evaluated across 20 independent simulation trials per cell for a unidirectionally coupled chaotic logistic map system ($N = 300$ observations). Grid axes span 11 dynamical coupling strengths ($C \in [0.00, 0.50]$) and 5 observational Gaussian noise levels $\sigma_{\text{noise}} \in [0.01, 0.20]$ expressed as a ratio of series standard deviation). Causal significance was determined using the analytical threshold ($T_{\text{crit\_TS}}$) at nominal $\alpha = 0.05$.

**(a) True causal direction ($X \rightarrow Y$).** In the uncoupled baseline ($C = 0.00$), the detection rate remained strictly 0.0% across all noise levels. Under weak forcing ($C = 0.05$), statistical power exhibited an expected gradual onset (15% to 30% under low-to-moderate noise). Once coupling reached $C \geq 0.15$, F-CODEC achieved and maintained 100% statistical power across all noise regimes, retaining full sensitivity even under severe 20% observational noise contamination ($\sigma_{\text{noise}} = 0.20$).

**(b) False causal direction ($Y \rightarrow X$).** In the uncoupled reverse direction, the detection rate remained precisely 0.0% across all 55 grid cells. Even under strong coupling regimes ($C \geq 0.35$) where conventional state-space reconstruction methods suffer from runaway false positives due to generalized synchronization, nonparametrically conditioning on the target's intrinsic history via adaptive forest kernels prevented spurious feedback detections.

### Supplementary Note 3: High-Throughput Scalability and Computational Runtime Benchmarks

A fundamental barrier to deploying causal inference in real-time streaming analytics, high-frequency IoT sensing, and genome-scale network discovery is computational costs. Conventional nonparametric frameworks—exemplified by Convergent Cross Mapping (CCM) and Transfer Entropy (TE)—rely heavily on surrogate testing, in which the observed index value (e.g., Pearson's correlation coefficient in CCM) calculated by a pair of original time series is compared with null distribution made by an original time series and hundreds of phase-randomized surrogate time series made from the opposite time series, to examine the statistical significance while preserving temporal autocorrelation. This surrogate testing process imposes severe computational costs, increasing with time-series length.

#### 3.1| Benchmarking Protocol and Experimental Setup

To evaluate the end-to-end execution efficiency of the full hypothesis-testing pipeline (from raw observational data ingestion to the final binary significance verdict at nominal $\alpha = 0.05$), we compared F-CODEC against standard implementations of CCM and information-theoretic causality across expanding sample lengths ($N \in \{100, 250, 500, 1000, 2000\}$ observations):

- **F-CODEC (Surrogate-Free):** Evaluated using Generalized Random Forests (grf, $B = 500$ trees) and the analytical time-series threshold $T_{\mathrm{crit\_TS}}$. Because the null threshold is evaluated analytically directly from the empirical rank projection and lag-1 autocorrelation, the entire test requires only a single pass of forest estimation without generating any surrogate series.

- **CCM (rEDM v0.7.5):** Evaluated using simplex projection across reconstructed delay manifolds ($E = 2$, $\tau = 1$, $tp = -1$). Statistical significance was assessed by benchmarking the observed cross-map skill ($\rho$) against an empirical null distribution derived from 100 Ebisuzaki phase-randomized surrogate time series.
- **Transfer Entropy via Unified Information Causality (rUIC):** Evaluated using state-space information transfer ($E = 2$, $tp = 1$), likewise assessed against 100 Ebisuzaki phase-randomized surrogates.
- **Hardware Standardization:** All benchmarks were timed using "tictoc" package on an isolated multi-core workstation CPU: Intel(R) Xeon W-2102 operating at 2.9 GHz with RAM 16 GB DDR4 memory running R version 4.3.2. Workloads were parallelized across 4 parallel workers via future::multisession.

#### 3.2| Runtime Scaling and Speedup Profiles

The empirical runtime trajectories demonstrate a decisive order-of-magnitude efficiency advantage for F-CODEC across all sample sizes (Supplementary Fig. S3).

**Figure S3**

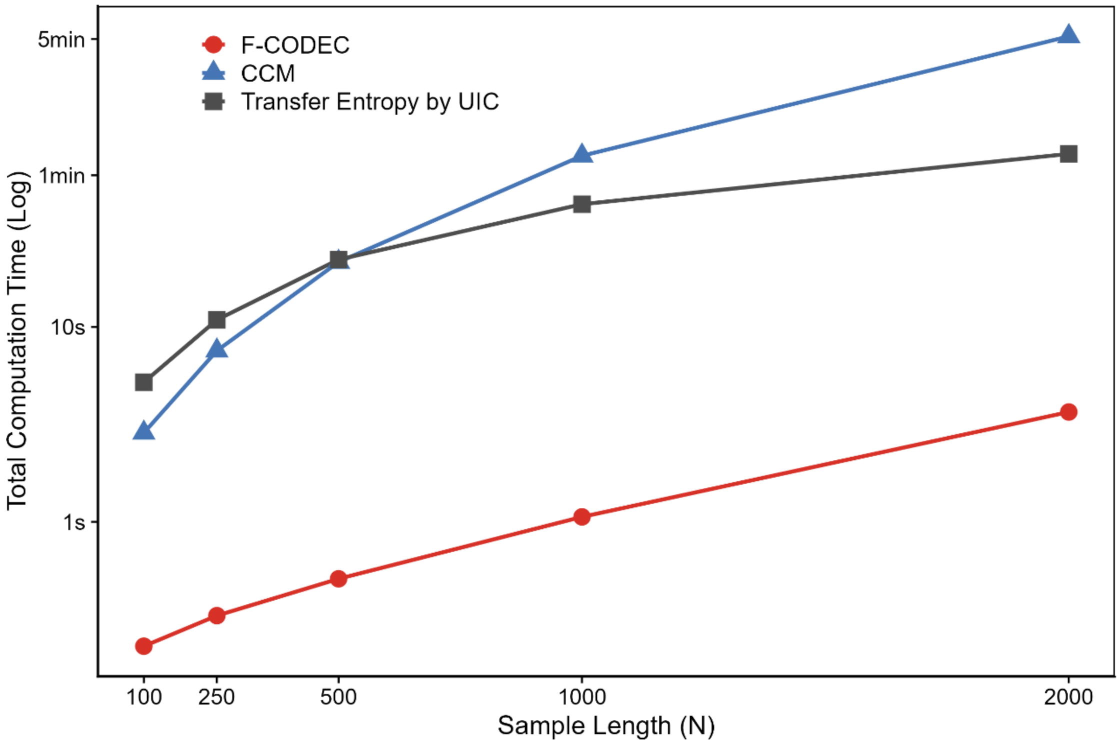


**Figure S3. End-to-end computational runtime comparison across time-series sample lengths.** Log-scale total execution time (seconds/minutes) required to complete a full bivariate causal hypothesis test (observed calculation and significance testing at nominal $\alpha = 0.05$) across sample lengths $N \in [100, 2000]$ by F-CODEC (red circles and solid line), CCM (blue triangles and solid line) and Transfer Entropy by UIC (dark grey squares and solid line). Benchmarks were executed on an identical multi-core computing platform.

F-CODEC completed the entire causal hypothesis test in 0.25 seconds for $N = 100$, 0.55 seconds for $N = 500$, and reached only 3.6 seconds at $N = 2000$. The execution time scaled smoothly with sample size, reflecting the efficient $O(B{\cdot}N\log N)$ computational complexity of honest regression forest construction. In contrast, surrogate-dependent approaches exhibited computational scaling. CCM required 3.0

seconds at $N = 100$, escalating to 25.4 seconds at $N = 500$, and reached 312.4 seconds (over 5.2 minutes) at $N = 2{,}000$. Similarly, Transfer Entropy via rUIC required 5.2 seconds at $N = 100$, 26.8 seconds at $N = 500$, and 85.6 seconds (1.4 minutes) at $N = 2000$. At $N = 2{,}000$, F-CODEC was approximately 87-fold faster than CCM and 24-fold faster than UIC-TE. If evaluated against the gold-standard 1,000-surrogate protocol recommended for high-precision null rejection, CCM would require nearly an hour (~52 minutes) per bivariate test at $N = 2{,}000$, whereas F-CODEC provides an immediate decision in 3.6 seconds. This dramatic reduction in computation time proves that F-CODEC removes the surrogate-data bottleneck, unlocking practical real-time causal tracking in continuous sensor streams and edge-computing microcontrollers.
